# Sub-Chronic Chlorpyrifos Exposure Leads to Epigenetic and Sex-Specific Behavioral Changes in Adult Mice

**DOI:** 10.64898/2026.06.08.730429

**Authors:** Alyssa R. Daniel, Noah Gernander, Sara Dodge, Cessily Hayes, Emma Simpson-Wade, Emese H. C. Kovács, Gibson Dowd, Jared M. McLendon, Benjamin Hing, Marie E. Gaine

**Affiliations:** Interdisciplinary Graduate Program in Human Toxicology, University of Iowa, Iowa City, IA, USA; Department of Pharmaceutical Sciences and Experimental Therapeutics, College of Pharmacy, University of Iowa, Iowa City, IA, USA; Department of Psychiatry, University of Iowa Carver College of Medicine, Iowa City, IA, USA; Iowa Neuroscience Institute, University of Iowa Carver College of Medicine, Iowa City, IA, USA; Department of Molecular Medicine, University of Iowa Carver College of Medicine, Iowa City, IA, USA; Department of Neuroscience and Pharmacology, University of Iowa Carver College of Medicine, Iowa City, IA, USA; Division of Cardiovascular Medicine, Department of Internal Medicine, University of Iowa Carver College of Medicine, Iowa City, IA, USA; Department of Molecular Physiology and Biophysics, University of Iowa Carver College of Medicine, Iowa City, IA, USA

**Keywords:** Organophosphate(s), Pesticides, Epigenetics, Estrogen, Sex-specific

## Abstract

Chlorpyrifos is a widely used organophosphate pesticide that exerts its primary toxic effect through inhibition of acetylcholinesterase (AChE). Although the acute neurotoxicity of chlorpyrifos is well characterized, the lasting biochemical, behavioral, and epigenetic consequences of sub-chronic exposure remain poorly understood, particularly when considering sex-specific differences. Therefore, we exposed male and female C57BL/6J mice to either peanut oil (n=19), low chlorpyrifos exposure (1 mg/kg/day; n=19), or high chlorpyrifos exposure (10 mg/kg/day n=10) repeatedly for 21 days via subcutaneous injection. Blood AChE activity, behavior, and hippocampal DNA methylation were measured across groups. During exposure, AChE activity decreased in both males and females but only returned to baseline after behavioral testing in females exposed to low chlorpyrifos levels. Behavioral tests also revealed a sex-specific phenotype, with females in the low exposure group exhibiting reduced forced swim test immobility and a significant time by exposure interaction in open field habituation. No significant behavioral effects were observed in males. Significant DNA methylation changes were observed at 3,538 CpG sites in male and female mice after high exposure. Sex-specific analyses revealed two female-specific differentially methylated CpGs after high exposure. Pathways enriched for differentially methylated genes included several related to synaptic remodeling, cholinergic synapse, and various endocrine systems. These findings demonstrate that repeated high chlorpyrifos exposure leads to persistent cholinergic disruption and DNA methylation changes. However, the female-specific behavioral changes seen are independent of AChE activity and widespread DNA methylation changes, suggesting additional mechanisms, present only in females, may underlie behavioral sensitivity to chlorpyrifos.

**New and Noteworthy:** Sub-chronic chlorpyrifos exposure in adult mice produced dose-dependent blood AChE suppression and widespread hippocampal DNA methylation changes in both sexes, with pathway enrichment including cholinergic synapse and endocrine systems. Behavioral effects were subtle, with females in the low exposure group showing reduced forced swim immobility and altered locomotor habituation. Notably, the female-specific behavioral changes seen are independent of AChE activity and DNA methylation changes, suggesting novel mechanisms may underlie female behavioral sensitivity to chlorpyrifos.

## Introduction

Chlorpyrifos has been one of the most widely used organophosphorus pesticides (OP) since 1965 due to its low cost and effectiveness against a broad range of insects [1, 2]. Exposure to chlorpyrifos can occur via occupational exposure as well as ingestion of treated fruits and vegetables [3]. Regulation of chlorpyrifos use varies around the world, with regulations debated in the United States as recently as 2023 [4], and current applications allowed for alfalfa, apple, asparagus, cherry (tart), citrus, cotton, peach, soybean, sugar beet, and wheat (spring and winter) produce [5]. Additionally, despite bans in other countries, persistent levels of chlorpyrifos are still being measured, as seen in Egyptian tomato paste [6] and bird feathers in the United Kingdom and Republic of Ireland [7]. Chlorpyrifos, and its primary metabolite chlorpyrifos-oxon, is absorbed into the blood and distributed throughout the body including within the liver, brain, kidneys, and fat stores [8]. Due to its lipophilicity, chlorpyrifos can cross the blood-brain barrier, which plays a major role in its contribution to neurological symptoms [9]. Chlorpyrifos is known to be acutely neurotoxic due to its ability to inhibit the acetylcholinesterase (AChE) enzyme, which leads to an overstimulation of the post-synaptic neuron because of an excess of acetylcholine [9]. Acute exposures are associated with cholinergic crisis affecting neurological, cardiovascular, and respiratory function [10], with high acute exposure linked to cholinergic syndrome, seizures, and death [11]. Health effects of low and moderate repeated occupational and environmental exposures to chlorpyrifos are not well understood mechanistically, but it has been suggested that cholinergic inhibition may not be responsible for low exposure effects [12, 13].

Long-term effects of chlorpyrifos exposure such as memory impairment, confusion, anxiety, personality changes, and peripheral neuropathy have been observed in humans and animal studies [14–19]. Importantly, chlorpyrifos exposure has also been associated with depression in several studies [20–22], with chronic low level exposure associated with neurobehavioral deficits [12, 18, 23–27]. A notable 2022 study by Ribeiro and colleagues showed that when rats were repeatedly exposed to occupationally relevant levels of chlorpyrifos for 21 days (5 or 10□mg/kg/day), depressive-like behavior was observable 11 weeks post-washout period, indicating that sub-chronic exposure can lead to depressive-like symptoms in animal models even after AChE rescue [17].

There are several noncholinergic mechanisms of chronic OP toxicity suggested in the literature including epigenetic mechanisms [28–31]. Differentially methylated patterns have been found after exposure to a range of OP [30, 32] and higher levels of a chlorpyrifos metabolite (3,5,6-TCP) have been linked to increased DNA methylation [28]. In addition, gene-specific DNA methylation changes have been associated with prenatal chlorpyrifos exposure in cord blood samples [33] and postnatal exposure in ApoE-TR mice [34]. At the mechanistic level, chronic chlorpyrifos exposure of a human liver cell line caused a reduction of DNMT3A and DNMT3B [35] and widespread DNA methylation changes [36]. Similarly, mice exposed to chlorpyrifos via oral gavage for 120 days showed a reduction in Dnmt1 in the liver and hepatic hypomethylation for low but not high exposure [37], and increased expression of Dnmt3a and Dnmt3b were found in ovarian samples of rats exposed to low levels of chlorpyrifos [38]. Even transgenerational DNA methylation and gene expression changes have been found in offspring after low chlorpyrifos exposure of pregnant mice [39, 40]. Together, these data suggest that chlorpyrifos directly impacts DNA methyltransferase expression leading to DNA methylation changes after chronic exposure.

In this study, we used low and high subcutaneous chlorpyrifos exposure to investigate the relationship between AChE inhibition, behavior, and hippocampal DNA methylation after sub-chronic exposure in adult male and female mice. We demonstrate that high exposure produces widespread hippocampal DNA methylation changes in both sexes, while subtle behavioral effects emerged exclusively in females following low exposure, indicating both sex-specific and dose-dependent effects.

## Materials and Methods

### Animals

All work was completed in compliance with the Office of the Institutional Animal Care and Use Committee (IACUC) guidelines in place at the University of Iowa, and all procedures were outlined in the lab’s approved animal protocol. Adult male and female C57BL/6J mice (3 months old; JAX:000664) were exposed to chlorpyrifos via subcutaneous injection for 21 continuous days. Chlorpyrifos (Chem Service Inc.) was dissolved in peanut oil (ThermoFisher) by vortexing. Stock solutions were stored at 4°C between dosages. The low exposure group received ∼1 mg/kg/day (n=9M, 10F) and the high exposure group received ∼10 mg/kg/day (n=5M, 5F). Exposure levels were determined by a literature search of occupational dosage levels in animal models of chlorpyrifos exposure [17, 41, 42]. Controls (n=9M, 10F) received vehicle peanut oil in direct relation to their body weight (but not exceeding 250 μL per IACUC guidelines). Mice were weighed every day to determine the volume of their next dose and to monitor signs of toxicity.

### Blood Collection and AChE Assay

Blood (∼100 µl) was collected from either via the tail or submandibular vein at Day 0 (one day before exposure began) and then collected every seven days (Days 7, 14, 21) including time of euthanasia (Day 37) to assess AChE activity as a biochemical marker of chlorpyrifos exposure. On each day of blood collection, a colorimetric AChE assay (SigmaAldrich) based on the Ellman method was conducted [43] with a BioTek Epoch microplate spectrophotometer (Agilent Technologies) set at 412 nm at 1-minute intervals for 5 minutes. Enzyme activity (Units/L) was determined by a kinetic curve, replicate results averaged and then normalized to initial concentration.

### Behavioral Testing

Mice were provided with food and water ad libitum with enriched paper bedding used in all cages. All experiments were conducted during the animals’ light cycle and animals were housed in groups of up to five. All mice were group housed. All equipment was cleaned between trials with 70% ethanol. Mice were acclimated to the Behavior Core (University of Iowa) on Days 22-23 for open field test (OFT; occurring on Day 25), tail suspension test (TST; occurring on Day 27), and forced swim test (FST; occurring on Day 29) and Days 30-32 for sucrose preference test (SPT; occurring on Days 33-37). Euthanasia occurred immediately after data collection on Day 37. During behavioral testing (16 days), the mice were not injected with vehicle or chlorpyrifos. Prior to behavioral testing, animals were acclimated to the behavior room for 30 minutes with testing carried out as follows:

#### OFT

Locomotion, habituation, and anxiety-related behavior were assessed using the OFT. Mice were put into a 40 cm x 40 cm arena and allowed to explore for 30 minutes. An overhead ring light was used to provide 115 lux across the arenas. Mice were recorded with an overhead camera using Pylon Viewer. Data analysis was conducted with EthoVision XT (Noldus) software as cumulative totals and in 10-minute time increments (0–10, 10–20, and 20–30 minutes).

#### TST

Escape-oriented behavior or lack thereof, represents a depressive-like phenotype in rodents, and was tested using the TST. As C57BL6/J mice are prone to tail-climbing behavior, a plastic tube was put over the tail to prevent this [44]. Mice were recorded for six minutes captured on video by Pylon Viewer (Basler). Behavioral Observation Research Interactive Software (BORIS v8.20.4) [45] was used to score all six minutes of each video. Mobility was defined as genuine mobility, involving moving all four limbs, violent shaking, reaching for the bar, or trying to climb [44]. As established by Can et al., movement that was confined to the front legs without the involvement of the hind legs, oscillations or pendulum swings due to momentum, or exhibiting a sitting position after climbing was not considered movement in accordance with established protocol [44].

#### FST

An additional test of depressive-like behavior was the FST. Mice were introduced to the water (20-25°C) gently to avoid submerging them under water, and recording was started and continued for six minutes. The video was monitored throughout recording to ensure mice did not dip below surface level. BORIS was used to score videos for mobility vs. immobility, with immobility defined as floating or making only the movements necessary to keep the head above water, and mobility defined as swimming or climbing-like behavior, in which mice aggressively pawed at the walls of the cylinder.

#### SPT

Anhedonia was assessed using the SPT. Briefly, rodents were provided two 25 mL serological pipettes modified with sipper tubes for drinking water over a two-day training period. During the testing period, rodents were given the opportunity to drink either from 2% sucrose or sucrose-free water over the span of four days. Pipettes were switched between left and right each day to negate side preference effects [46]. Volumes were measured twice a day, and sippers were filled when necessary.

### DNA Methylation Analysis

After the behavioral tests on Day 37, the hippocampus was collected immediately after cervical dislocation, flash-frozen, and kept at -80°C until DNA extraction. DNA was extracted from brain tissue with a DNeasy Blood and Tissue Kit (Qiagen) with RNase treatment. DNA concentration was calculated, and purity was confirmed by NanoDrop One (ThermoFisher).

Infinium Mouse Methylation Beadchip Arrays (Illumina) were used to measure hippocampal DNA methylation. Genomic DNA (500 ng) was bisulfite converted using an EZ DNA Methylation Kit (Zymo Research). Samples were then prepared for hybridization according to manufacturer’s instructions. Arrays were analyzed by the Iowa Institute of Human Genetics (University of Iowa).

BeadChip array data was pre-processed using the *SeSAMe* (Sensible Step-wise Analysis of DNA Methylation BeadChips, version 1.24.0) pipeline in R Studio (version 4.4.1) [47]. Pre-processing included strain inference, masking probes of poor quality, channel inference, dye bias correction, detection p-value calculation, and background subtraction. After data was normalized to match Infinium I/II probe distribution, beta-values were retrieved, the final matrix of normalized beta-values was further filtered by removal of non-CpG or low-quality probes and any probes that were missing in more than 5% of samples. Filtered beta-values associated with 265,284 CpG sites were ultimately used in differentially methylated locus (DML) analysis using the DML function in *SeSAMe*. DMLs were modeled by exposure and sex was controlled for as a covariate. False Discovery Rate (FDR) corrected P-values were calculated by the *p.adjust* function with the “fdr” option at a significance threshold of 0.05. The FDR-adjusted results were annotated with genes from mm10 using *annotatr* (v1.28.0). Annotation tracks were first generated using the *build_annotations* function with argument “genome = mm10” and overlapped with probe coordinates using *annotate_regions* function. This information was then merged with the DML results based on probe ID. A KEGG pathway enrichment analysis was run with the annotated gene list using the *enrichKEGG* function in the *clusterProfiler* (version 4.10.0) package with the mouse annotation package *org.Mm.eg.db* (v3.18.0). A secondary analysis to investigate sex-specific findings was completed by re-analyzing the beta-values from males and females separately using the DML function modeled by exposure.

### Statistical Analyses

GraphPad Prism (version 10.6.1) was used to conduct statistical testing of all enzymatic assay and behavior data. For recorded behavior, the average between two scorers was used, and inter-scorer reliability was measured as a relative standard deviation less than 20%. If above this relative standard deviation, the videos were re-scored by both individuals. Average values were used in statistical analyses. Blood AChE activity, OFT 10-minute increment travel distance, SPT data, weekly average body weight, and OFT 10-minute increment inner zone time were analyzed using a mixed-effects models with Geisser-Greenhouse correction and either Dunnett’s or Tukey’s correction for multiple testing. This approach was selected over traditional repeated-measures ANOVA as it does not assume sphericity and accommodates missing data. Prior to analysis, SPT values were arcsine square root transformed to account for the bounded nature of percentage data and the ceiling effect observed in the raw values. Results are expressed as original SPT percentages for interpretability, with statistical analyses performed on transformed values. Cumulative OFT travel distance, cumulative OFT inner zone time, FST immobility, and TST immobility were analyzed using one-way ANOVA with Brown-Forsythe test for equality of variances. Where variances were found to be unequal (FST and TST), Welch’s ANOVA with Dunnett’s correction for multiple testing was applied. Males and females were analyzed separately throughout, except for the main DNA methylation analysis. Statistical significance was defined as P<0.05.

## Results

### Weight and Physical Observations

An expected increase in weight over time was noted in both males and females (P<0.0001) but no significant weight changes (P>0.05) were seen across the exposure groups at each time point (**Supplemental Fig. S1**). No physical symptoms of cholinergic crisis such as salivation or lacrimation were observed during the experimental period.

### Blood Plasma AChE Assay

After both low and high chlorpyrifos exposure, normalized AChE activity was significantly reduced in both males and females at Day 7, 14, and 21 (all P<0.005; **Fig. 1**). In males, reduced activity was also seen at Day 37, 16 days after exposure ended, in the low exposure (mean difference = 40.05, 95% CI [20.02, 60.09], P=0.0006) and high exposure (mean difference = 85.59, 95% CI [65.86, 105.3], P<0.0001) groups. Notably, in females, there was lower activity with low chlorpyrifos exposure at Day 14 and 21 compared to males (all P<0.0001), however this activity returned to baseline by Day 37 (mean difference = 13.92, 95% CI [-7.900, 35.74], P=0.2350). Continued suppression was seen in the female high exposure group at Day 37 (mean difference = 80.32, 95% CI [58.90, 101.7], P<0.0001).

**Figure 1.**
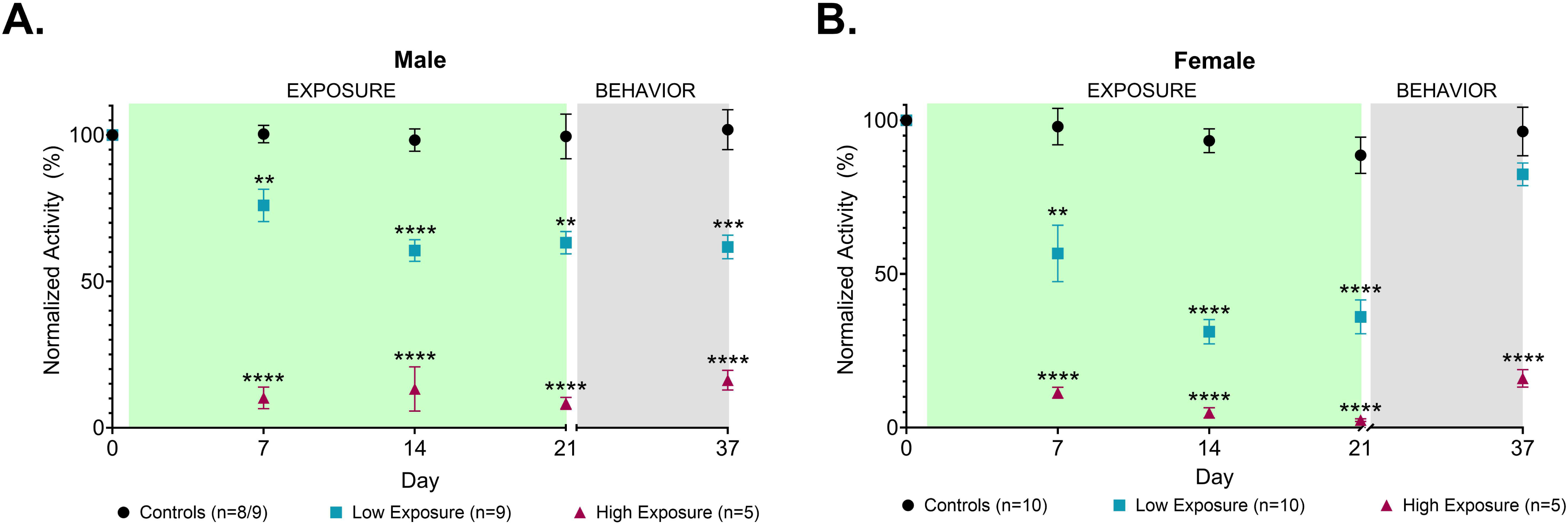
Chlorpyrifos exposure causes sex-specific changes to acetylcholinesterase (AChE) activity over time. **A.** In male mice, after 21 days of low or high chlorpyrifos exposure there was a significant reduction in normalized AChE activity (P<0.0001). This reduction remains significant even 16 days after exposure ended (low P=0.0006; high P<0.0001). **B.** In female mice, after 21 days of low or high chlorpyrifos exposure there was a significant reduction in normalized AChE activity (P<0.0001). However, after low exposure, AChE activity returned to baseline by Day 37 (P=0.2350). One male control mouse was excluded on Day 37 due to missing data. Data are expressed as mean□±□s.e.m. ** denotes P<0.005; *** denotes P=0.0006; **** denotes P<0.0001

### Behavioral Results

OFT provided data on locomotion, habituation, and anxiety-like behavior (**Fig. 2**). In males, there was no significant difference across exposure groups when assessing distance traveled as a cumulative value or in 10-minute increments. In females, a similar trend was found with no significant difference seen in cumulative distance traveled. However, when comparing 10-minute increments, a significant time by exposure interaction was detected (F (2.812, 29.53) = 4.314, P=0.0137). However, post-hoc pairwise comparisons using Tukey’s test did not reveal significant differences between any exposure groups at any individual time increments (lowest P=0.0661). There was also no significant difference in time spent in the inner area during OFT (**Supplemental Fig. S2**).

**Figure 2.**
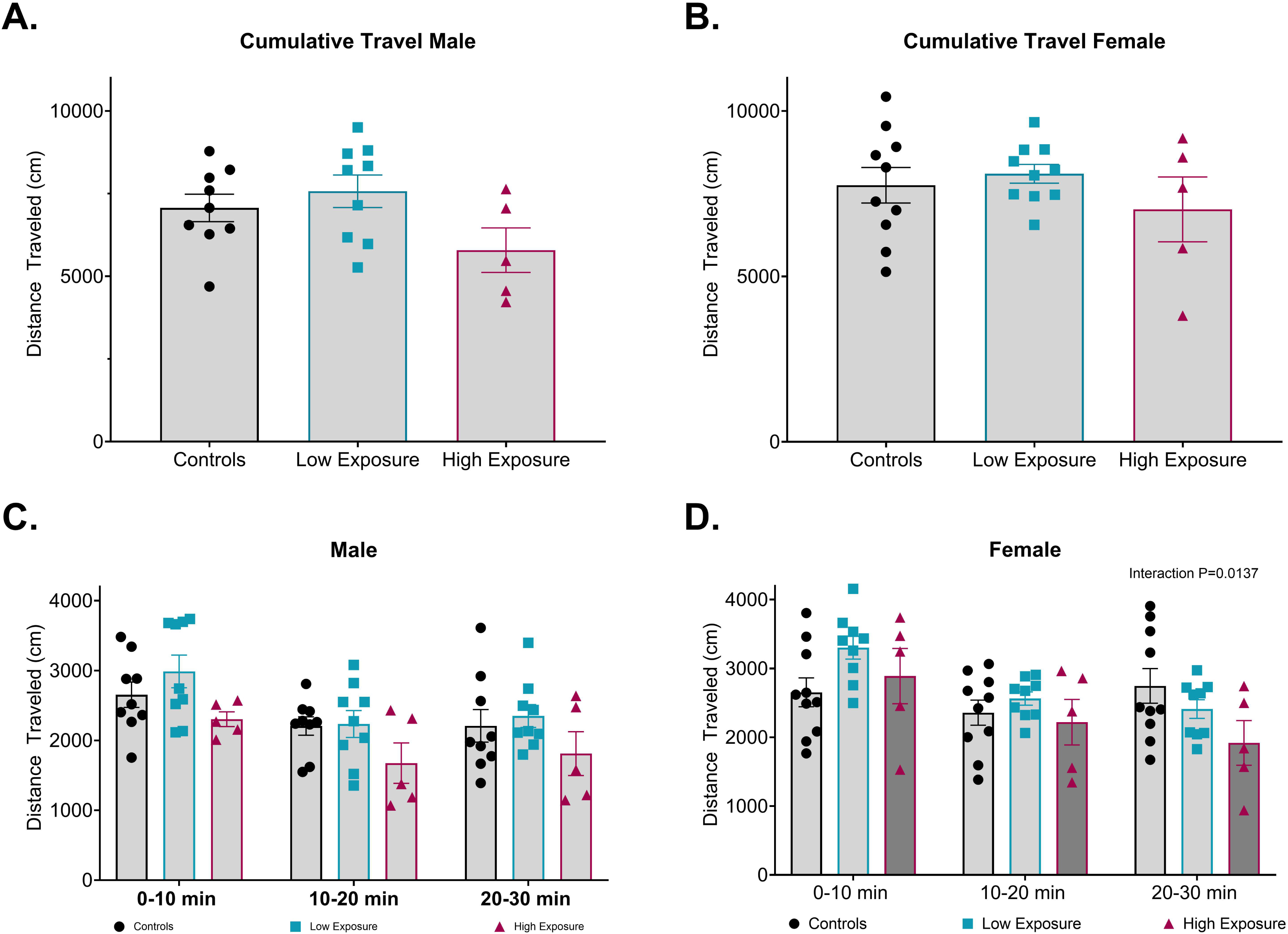
Chlorpyrifos exposure causes a female-specific locomotor habituation using the Open Field Test (OFT). **A.** and **B.** After 21 days of low or high chlorpyrifos exposure in male and female mice there was no difference in cumulative distance travelled in the OFT. **C.** When distance traveled is separated into 10-minute increments, there is no difference in male mice but **D.** in female mice, we see a significant Time x Exposure interaction between time and exposure (P=0.0137). Post-hoc pairwise comparisons using Tukey’s test did not reveal significant differences between any exposure groups at any individual time increments (lowest P=0.0661). One female mouse in the low exposure group was removed due to missing data. Data are expressed as mean□±□s.e.m.

The TST and SPT assessed depressive-like behavior and no significant differences were observed in males or females (**Fig. 3**). Similarly, there was no significant difference in immobility in the FST when analyzing the male groups. However, Welch’s ANOVA revealed a significant effect of exposure group on immobility time in females (W (2.00, 11.44) = 4.479, P=0.0366; **Fig. 3D**). Post-hoc pairwise comparisons using Dunnett’s test showed that females exposed to low chlorpyrifos levels had less immobility duration (128.6 seconds) than controls (164 seconds; mean difference = 35.42, 95% CI [2.267, 68.58], P=0.0359).

**Figure 3.**
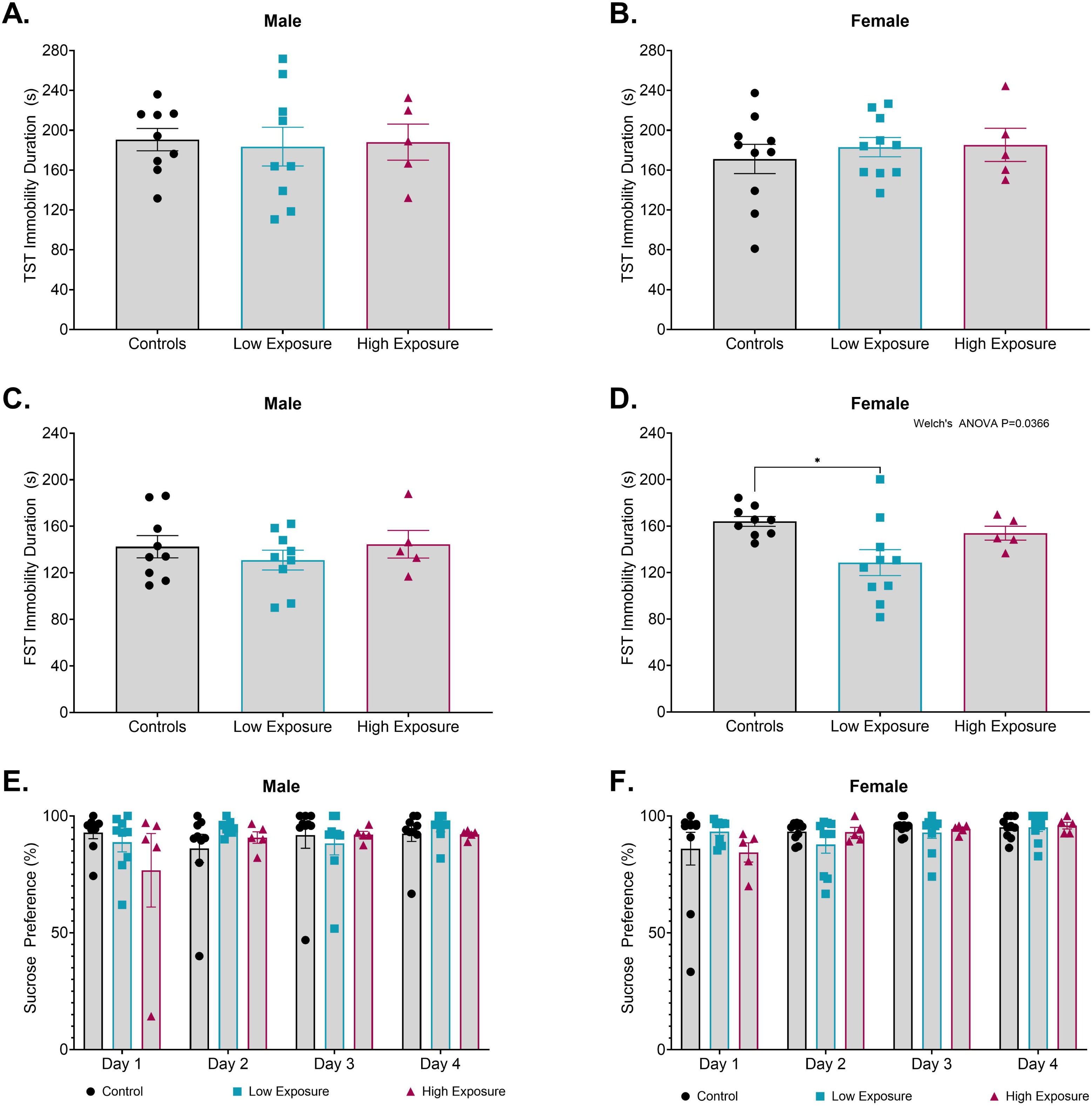
Chlorpyrifos exposure induces subtle behavioral changes in female mice. **A.** and **B.** No significant difference in immobility duration was noted in the Tail Suspension Test (TST). **C.** No significant difference in immobility during the Forced Swim Test (FST) was found in male mice but **D.** in female mice a significant effect of exposure group on immobility time (Welch’s ANOVA P=0.0366) was seen along with a significant decrease in immobility in female mice after low chlorpyrifos exposure (P=0.0359). **E.** and **F.** No significant difference in preference percentage was noted in the Sucrose Preference Test. One control female mouse in the FST test data was excluded due to missing data. Data are expressed as mean□±□s.e.m. * denotes P<0.05

### Differential Methylation Results

After FDR correction and controlling for sex, no CpG sites were significantly differentially methylated after low chlorpyrifos exposure (FDR=0.999). A total of 3,538 CpG sites were significantly differentially methylated between the high exposure group and controls (FDR<0.05; **Supplemental Table S1** and **Fig. 4**). For the high exposure group, we also conducted a sex-specific comparison of differentially methylated CpG sites and no significant results were observed in males. However, two CpG sites were significantly differentially methylated when comparing controls to females in the high exposure group (**Supplemental Figure S3**). They were: cg43966652 (FDR=0.0440) annotated to Zinc finger protein 384 (*Zfp384)*, and cg46968954 (FDR=0.0440) annotated to Myosin VC (*Myo5c*).

**Figure 4.**
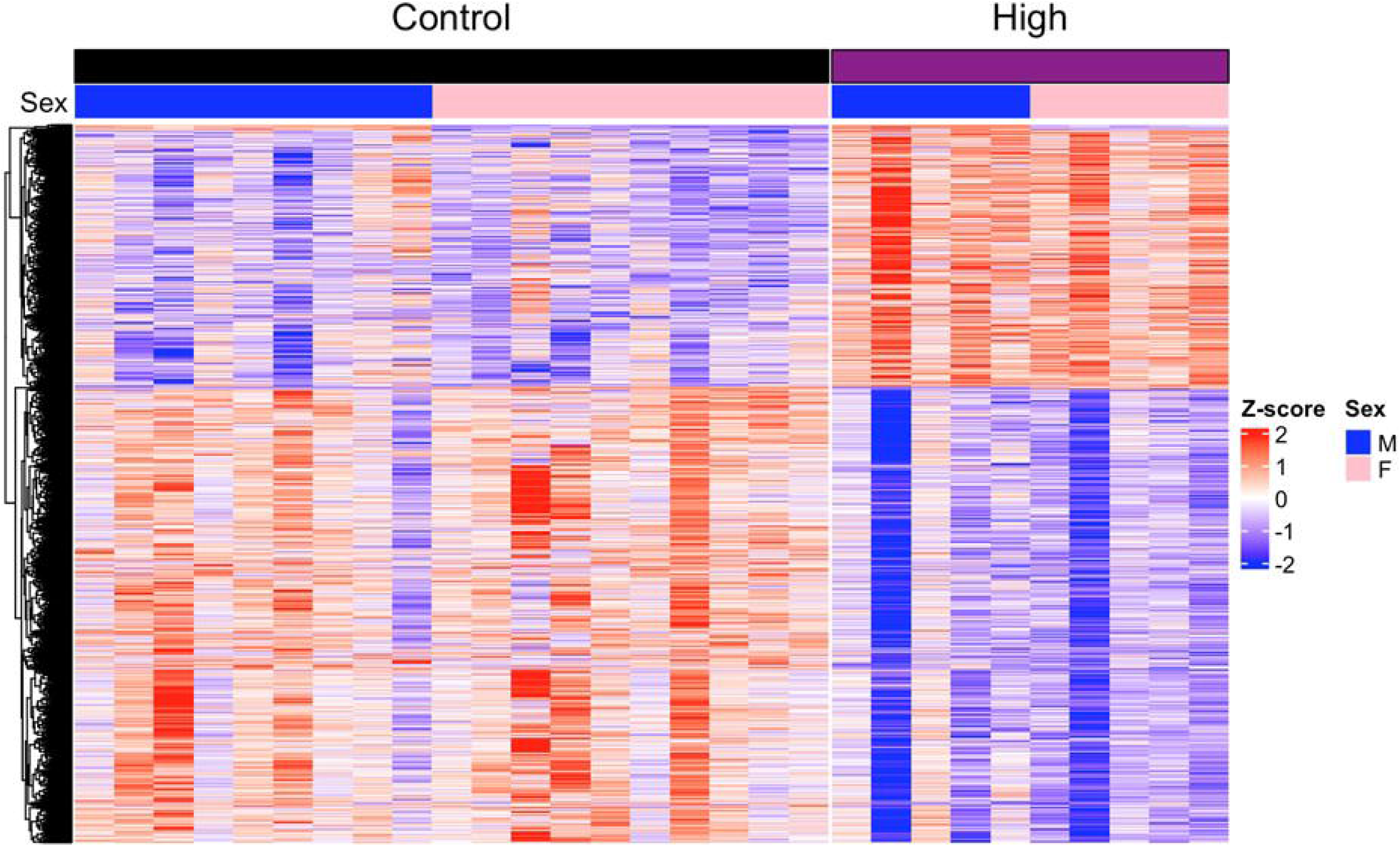
High chlorpyrifos exposure causes differentially methylated CpG sites in the hippocampus. Significant beta-value levels (FDR<0.05) are presented in a heat map for the control and high exposure group (both sexes). Low chlorpyrifos exposure did not cause significant changes in DNA methylation. Sex is labelled with a horizontal bar at the top, where blue denotes male and pink denotes female.

The CpG sites associated with high exposure were annotated to genes and analyzed in a KEGG pathway enrichment analysis and 98 pathways were found to be significantly enriched (**Supplemental Table S2** and **Fig. 5**). Notable pathways include glutamatergic synapse (Q=2.58×10^-5^), cholinergic synapse (Q=6.32×10^-6^), and estrogen signaling pathway (Q=4.81×10^-4^).

**Figure 5.**
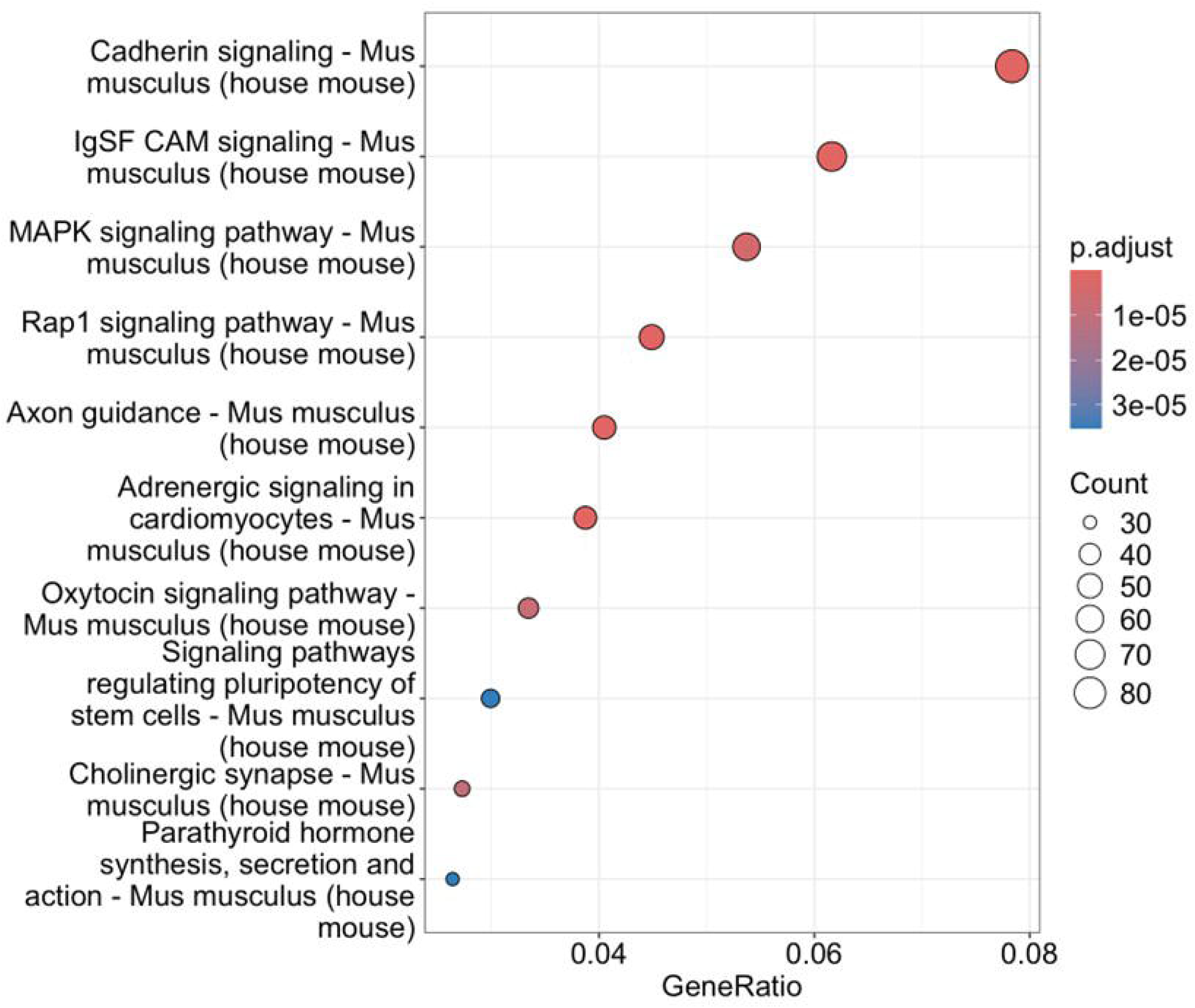
High chlorpyrifos exposure leads to differentially methylated genes enriched for specific KEGG pathways. This dot plot shows the top ten KEGG pathways, ordered by gene count, GeneRatio, and FDR adjusted P-value.

## Discussion

This study investigated the biochemical, behavioral, and DNA methylation effects of an occupationally relevant sub-chronic exposure to chlorpyrifos in adult male and female mice. Chlorpyrifos exposure produced dose-dependent suppression of blood AChE activity in both sexes throughout the exposure period, confirming effective biochemical disruption at both doses. When comparing males and females during exposure, a more dramatic reduction in activity was seen in females, which is consistent with human literature demonstrating elevated susceptibility to AChE inhibition in females after OP exposure [48–52]. However, the most notable finding at the biochemical level was the divergent recovery pattern between sexes in the low exposure group: females recovered to control levels by Day 37, while males remained significantly suppressed at the same timepoint. The novel finding of faster biochemical recovery in females may reflect hormonal differences, as estradiol modulates AChE activity [53].

Despite robust and persistent AChE suppression at both doses before behavioral testing, chlorpyrifos exposure did not produce significant changes in gross locomotor activity, anxiety-like behavior, or depressive-like behavior in either sex. This is in contrast with other studies observing depressive-like behavior in rodents after extended (≥10 days) exposure to chlorpyrifos, including TST, FST, and SPT [17, 54–57]. It is possible that depressive-like behavior was not seen in our study because there was not enough time between exposure and behavioral tests for the behavior to develop [17], or because our chlorpyrifos exposure paradigm increased vulnerability to subsequent stress, or stress sensitization, that requires a secondary stressor to unmask the expected behaviors [58]. Mode of exposure may also play a role in the differing results. We used subcutaneous injection because it has been shown to be useful in occupational models of dermal exposure in animal studies and results in a slower sustained release of the pesticide into circulation [59]. However, exposure route and vehicle of administration in chlorpyrifos experiments impacts metabolism and toxic response [59]. Despite the absence of expected behavioral changes across exposure groups, subtle but significant behavioral differences were detected exclusively in female mice. In the FST, females exposed to low chlorpyrifos levels moved more than controls. Reduced immobility in the FST could be interpreted as hyperactivity or increased fear/anxiety [60]. However, cumulative distance travelled and time in the inner area in the OFT were not different, arguing against the interpretation that reduced FST immobility reflects non-specific hyperactivity or anxiety in this case.

The significant time by exposure interaction in OFT locomotor habituation, while not resolved to specific pairwise comparisons after correction for multiple testing, reveals that chlorpyrifos exposed females differ from controls in the temporal pattern of activity across the 30-minute test session. These data suggest females in the high exposure group have a delayed habituation profile, with their activity declining in the 20-30 minute window rather than plateauing earlier as seen in controls and mice in the low exposure groups. Female mice exposed to chlorpyrifos displaying greater habituation time to a new environment may suggest decreased cognitive function or, together with the FST finding, may suggest that chlorpyrifos exposure in females leads to an absence of coping in certain stressful environments [61].

No behavioral effects were observed in males across any test. This is particularly striking given that males showed persistent AChE suppression at both doses through Day 37 and equivalent hippocampal DNA methylation changes compared to females. The absence of behavioral effects in males despite similar molecular disruption suggests that males are not simply unexposed to the consequences of chlorpyrifos, but they appear behaviorally resilient under the conditions tested, which has been observed elsewhere [62]. This is in contrast with human male-only studies that associated behavioral differences, including depressive symptoms, with chlorpyrifos exposure [63, 64].

Hippocampal DNA methylation changes, controlling for sex, were noted 16 days after high exposure ended. The enrichment of the cholinergic synapse pathway among differentially methylated genes provides a direct epigenetic correlate of the biochemical AChE suppression observed in this study and suggests that high chlorpyrifos exposure produces lasting molecular changes at genes central to cholinergic neurotransmission. The enrichment of multiple neurotransmitter systems, including glutamatergic, GABAergic, dopaminergic, and serotonergic synapse pathways, indicate that the epigenetic remodeling induced by high chlorpyrifos exposure extends beyond cholinergic circuitry to affect synaptic signaling broadly across the hippocampus [65–70]. Additionally, enrichment of several endocrine signaling pathways including thyroid hormone signaling, GnRH signaling, GnRH secretion, and estrogen signaling implicates the endocrine system in the epigenetic response to chlorpyrifos. The enrichment of estrogen signaling in particular is mechanistically relevant and is discussed below in the context of the female-specific behavioral findings. The female-specific DNA methylation findings, albeit small in numbers, also provide evidence of a female-specific epigenetic mechanism.

The broad methylomic remodeling of the hippocampus following high chlorpyrifos exposure is equivalent between sexes and, therefore, cannot account for the sex-specific behavioral outcomes. This divergence in results suggests a female-specific mechanism that, in combination with chlorpyrifos-induced epigenetic and biochemical disruption, produces the behavioral phenotype observed in females. One potential mechanism is estrogen as it modulates hippocampal cholinergic signaling at multiple levels, including AChE expression, cholinergic receptor density, and downstream intracellular signaling cascades [71], and regulates hypothalamic-pituitary-adrenal (HPA) axis activity, influencing the magnitude of stress responses and behavioral outcomes in assays such as the FST [72]. Chlorpyrifos is a known endocrine disruptor by impacting hormone biosynthesis and metabolism pathways [73] and disrupts the estrous cycle in rats [74]. Notably, estrogen signaling was a significantly enriched KEGG pathway in the hippocampal DNA methylation analysis, providing evidence from this study that chlorpyrifos exposure produces epigenetic changes at estrogen-regulated genes. Independent studies further support that chlorpyrifos disrupts estrogen biology, showing that chlorpyrifos metabolite levels are dose-dependently associated with decreased estradiol in human urine samples [75] and exposed fish [76], chlorpyrifos inhibits hepatic estradiol metabolism in humans [77], and chlorpyrifos exposure increases *ESR2* DNA methylation and decreases its gene expression in rat ovarian tissue [38]. There is also structural evidence that chlorpyrifos interacts directly with the human estrogen receptors disrupting estrogen signaling [78]. Recently, Pastorino et al., also showed that high chronic exposure to chlorpyrifos (10mg/kg/day from conception to six months old) in male mice selectively upregulated estrogen receptor beta but not estrogen receptor alpha and an elevated ratio of these receptors in the hypothalamus [79]. Together, these findings suggest that the endogenous estrogenic environment unique to females renders them behaviorally vulnerable to chlorpyrifos-induced disruption of estrogen-regulated hippocampal pathways as this interaction is absent in males. Estrogen levels and estrous cycle stage were not measured in the present study, and future work should directly test this mechanism.

### Limitations

Several limitations of the present study should be noted. The high exposure group was smaller than the control and low exposure groups (n=5 per sex), which reduced statistical power for detecting exposure effects, particularly in the sex-specific DNA methylation analysis. This epigenetic analysis was restricted to hippocampal tissue but sex-specific DNA methylation changes relevant to the behavioral outcomes may be present in other brain regions, such as the prefrontal cortex [80, 81]. DNA methylation was assessed at a single post-exposure timepoint, precluding conclusions about longitudinal patterns or whether the changes observed represent a stable or transient alteration. Estrogen levels and estrous cycle stage were not measured, which constrains the mechanistic interpretation of the sex-specific behavioral findings. The subcutaneous route of exposure, while appropriate for modeling occupational dermal absorption, was associated with oil deposit formation in most mice across all groups; although prior work suggests this does not affect behavioral outcomes in rats [82], it may introduce biological variability that should be addressed in future studies with larger sample sizes. Finally, behavioral testing was conducted at a single post-exposure timepoint (after 16 days), but the Ribeiro et al. study observed depressive-like behavior 11 weeks post-washout [17], suggesting that the full behavioral consequences of chlorpyrifos exposure may not yet have been established in the present paradigm.

### Conclusions

These findings contribute to a growing body of evidence that chlorpyrifos exposure in adulthood produces lasting molecular and behavioral effects at doses below those associated with overt toxicity and highlight the importance of considering sex as a biological variable in neurotoxicology research. The female-specific behavioral phenotype, reduced FST immobility and delayed OFT habituation, emerged during a period of recovering cholinergic inhibition and without female-specific DNA methylation changes, suggesting additional mechanisms that warrant further investigation. Future studies should assess estrogen levels and estrous cycle stage, measure gene expression changes associated with the identified DNA methylation changes, extend the analysis to additional brain regions, and examine behavioral outcomes at extended post-exposure timepoints to more fully characterize the trajectory and mechanisms of chlorpyrifos-induced neurobehavioral changes.

## Data Availability Statement

Data available on request from the corresponding author.

## Acknowledgements.

We would like to thank the Neural Circuits and Behavioral Core (NCBC) and Shane Heiney for help with the behavior analyses, and the Iowa Institute of Human Genetics (IIHG) for help with the DNA methylation analyses.

## Author Contributions

**ARD:** Conceptualization, Investigation, Methodology, Formal Analysis, Software, Writing-original draft, Visualization **NG:** Methodology **SD:** Methodology and Formal Analysis **CH:** Methodology **ESW:** Methodology and Formal Analysis **GD:** Formal Analysis **EK:** Formal Analysis **JMM:** Conceptualization, Methodology **BH:** Formal Analysis, Writing-original draft, Visualization **MEG:** Conceptualization, Supervision, Funding Acquisition, Writing-review and editing, original draft. All authors edited the manuscript.

## Supplemental Information

**Supplemental Figure S1.**
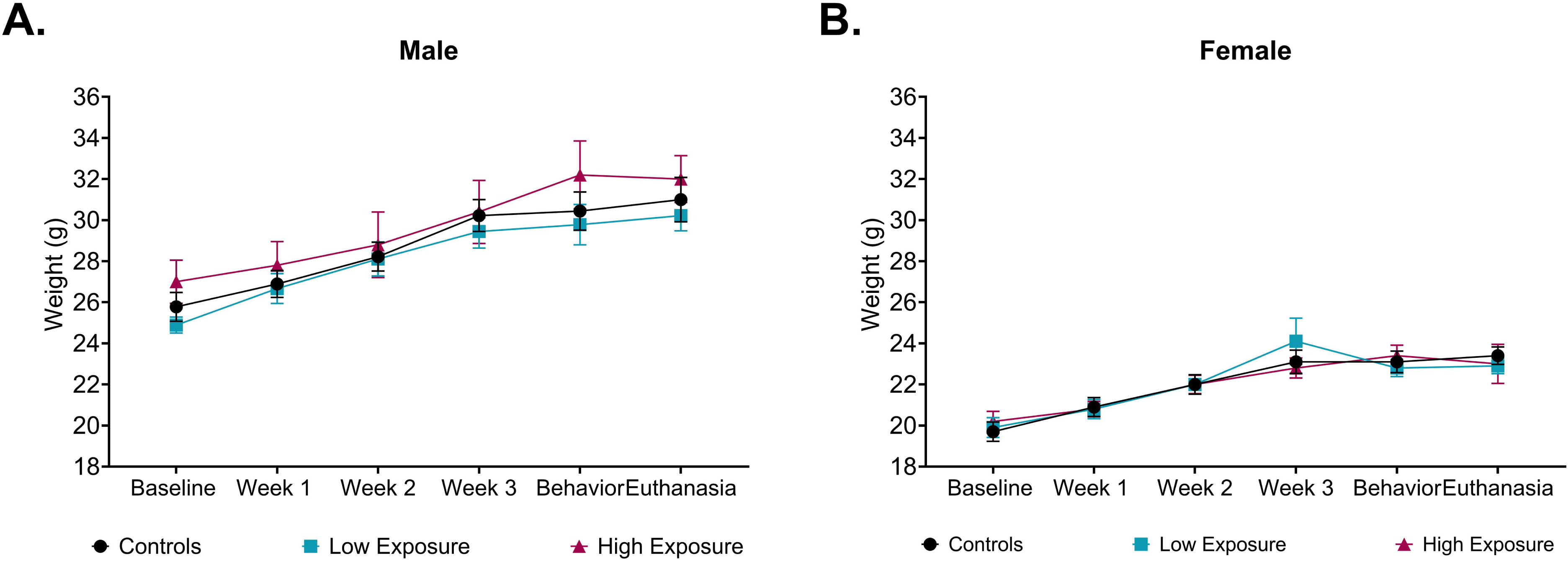
Chlorpyrifos exposure did not cause significant weight changes in **A.** male or **B.** female mice. Data are expressed as mean□±□s.e.m.

**Supplemental Figure S2.**
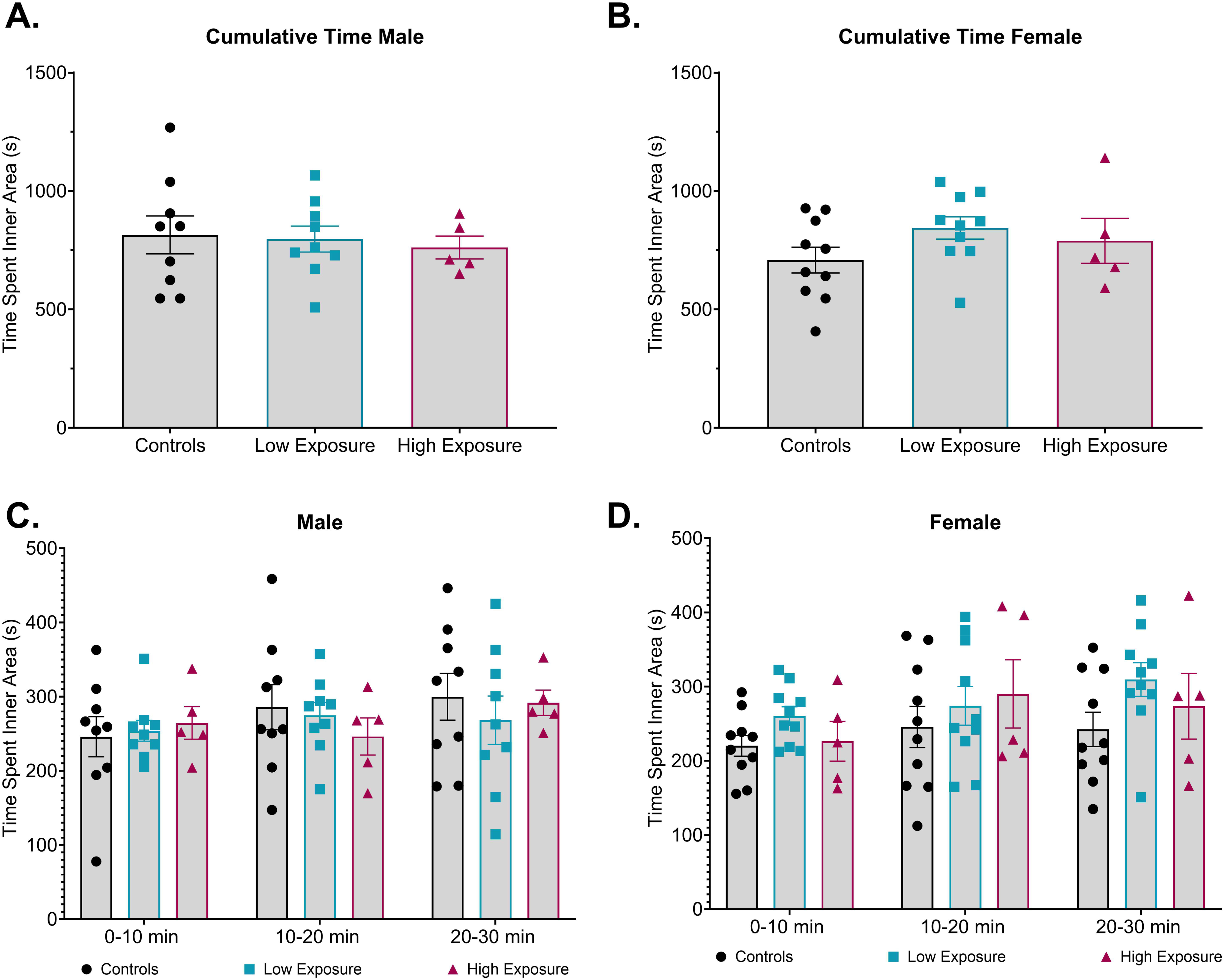
Chlorpyrifos exposure does not induce anxiety-like behavior in male or female mice. After 21 days of low or high chlorpyrifos exposure there was no difference in cumulative time spent in the inner area during the Open Field Test (OFT) in **A.** male or **B.** female mice. There was also no significant difference when time was separated into 10-minute increments in **C.** male or **D.** female mice. Data are expressed as mean□±□s.e.m.

**Supplemental Figure S3.**
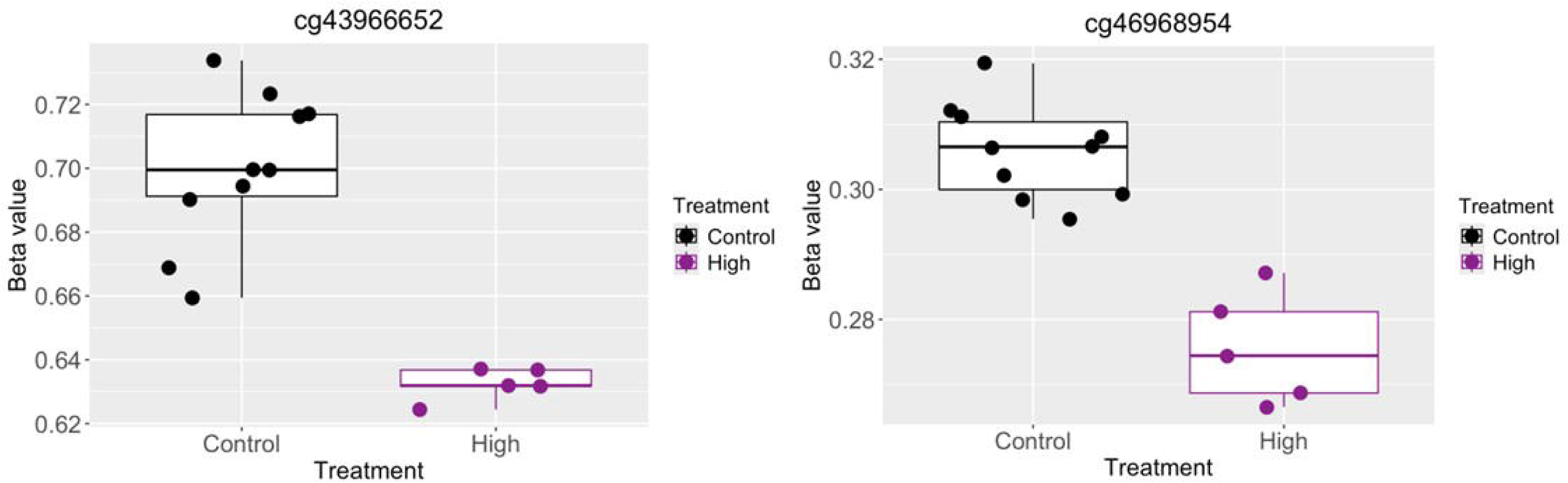
In sex-specific analyses, two CpG sites were significantly differentially methylated when comparing females in the high exposed group to female controls (FDR=0.0440 for both). **Left.** Cg43966652 is located in zfp384 (chr6:125035245; mm10). **Right.** cg46968954 is located in Myo5c (chr9:75244219; mm10). There were no male-specific significantly differentially methylated CpG sites.

**Supplemental Table S1.** Significantly Differentially Methylated CpGs After High Chlorpyrifos Exposure

**Supplemental Table S2.** Significantly Enriched Pathways after High Chlorpyrifos Exposure

